# Modeling the impact of genotype, age, sex, and continuous navigation on pathway integration performance

**DOI:** 10.1101/2023.09.11.556925

**Authors:** Dorothea Pink, Ekin Ilkel, Varnan Chandreswaran, Dirk Moser, Stephan Getzmann, Gajewski Patrick, Nikolai Axmacher, Hui Zhang

**Affiliations:** Department of Neuropsychology, Ruhr University Bochum, Germany; Department of Genetic Psychology, Ruhr University Bochum, Germany; Leibniz Research Centre for Working Environment and Human Factors, Germany

## Abstract

The Path Integration (PI) task stands as a fundamental activity within spatial navigation, encompassing a multifaceted underlying process that requires extensive computation of both translational and directional information during navigation. This task is postulated to engage the entorhinal cortex (EC), a critical brain region. Although rodent studies have firmly linked the EC with the PI task, such a correlation has yet to be definitively established in humans. Individuals harboring high-risk genes associated with Alzheimer’s disease (AD) present a unique opportunity to explore PI task performance in humans because EC is the first targeted region during the pathological development of AD. Furthermore, the execution of the PI task is also susceptible to the influences of age and gender. In order to methodically probe the influences of various factors on PI performance, we conducted a PI task in a cohort characterized by high-risk AD-associated genes in a virtual island environment with surrounding landmarks. Our findings unveiled intricate interactions between these risk genes, age, and gender in shaping PI performance, an influence primarily mediated through distance and angle estimation. This study provides an illuminating perspective on the investigation into the functional role of the entorhinal cortex within the context of the PI task.

## Introduction

Homing behavior entails the capacity to navigate directly back to a previously visited spatial location following an exploratory journey. This skill enables individuals to establish and maintain the connection between their current position and the reference point of their home base. While it constitutes a fundamental aspect of our daily activities, the process itself involves intricate computations of translational and directional parameters (Benhamou & Séguinot, 1995; Fujita et al., 1990; Harootonian et al., 2020; Klatzky et al., 1990; Maurer & Séguinot, 1995). To unravel the mechanisms behind homing behavior, researchers have designed the path integration (PI) task, a simulated version of this phenomenon (Klatzky et al., 1990; Loomis et al., 1993; Mittelstaedt & Mittelstaedt, 1980). During a path integration (PI) task, participants are initially directed to travel from a designated starting point to a specific target location. Subsequently, they are tasked with retracing their path back to the initial starting point along the shortest possible route. The PI performance is assumed to decline due to the accumulation of errors along navigation (Lakshminarasimhan et al., 2018; Lappe et al., 2007, 2011; Stangl et al., 2020). This intricate cognitive process has been attributed to the entorhinal cortex (EC), particularly due to the distinct properties of grid cells within it (Hafting et al., 2005; Moser & Moser, 2008). Grid cells, found in layers II and III of the medial entorhinal cortex (MEC), exhibit firing patterns at multiple locations forming grid-like configurations that span the entire spatial environment, hence their name. The regular distribution of firing spots enables grid cells to effectively monitor the distance between different grid positions, mirroring the demands of the PI task. Indeed, research involving genetic manipulation of grid cells in rodents has established their causal role in PI tasks (Gil et al., 2018; Tennant et al., 2018) . Despite the compelling evidence from rodent studies, a gap persists in our understanding of how dysfunction within the entorhinal cortex impacts path integration performance in humans.

Alzheimer’s Disease (AD), the predominant cause of dementia affecting 55 million individuals globally in 2019, is a progressive neurodegenerative disorder. The initial focal points of AD pathology are found within EC and the hippocampal gyrus (Braak et al., 1993). A significant genetic risk factor for the common late-onset form of AD is the ɛ4 allele of the Apolipoprotein E (APOE) gene (Bu, 2009; Farrer et al., 1997; Huang & Mucke, 2012) . Those possessing a single APOE4 allele have a 3-4 times heightened susceptibility to AD compared to those lacking this allele (Bertram & Tanzi, 2008; Corder et al., 1993) . AD symptoms manifest gradually, intensify over time, and eventually interfere with daily functioning. By studying individuals harboring high-risk genes of developing AD, we gained insights into how EC, a region implicated in AD, influences performance in PI tasks. Notably, young APOE4 carriers navigating a virtual environment displayed compromised grid-cell-like representations (Kunz et al., 2015). Recent findings indicate that healthy APOE4 carriers performed less accurately in pure PI tasks and angle estimation tasks compared to a control group (Bierbrauer et al., 2020; Coughlan et al., 2023). However, the precise connection between the malfunctioning entorhinal cortex and PI performance remains elusive (for an overview, refer to (Segen et al., 2022)).

Apart from the potential influence of pathological AD development on PI performance, this ability is also closely related to age and gender. Research consistently indicates that older individuals, when compared to their younger counterparts, exhibit more significant errors when performing the PI task (Adamo et al., 2012; Harris & Wolbers, 2012; Stangl et al., 2020) . The impact of gender on PI performance, however, presents a complex scenario (Fernandez-Baizan et al., 2019; Fortenbaugh et al., 2007; Goeke et al., 2013), often interacting with age (Yu et al., 2021) . Evidence has shown that morphological connections between the entorhinal cortex and other regions, such as the caudal anterior cingulate cortex and transverse temporal cortex, vary between male and female subjects (Nebli & Rekik, 2020) . Moreover, aging and AD neuropathology exert distinct effects on the physiology of entorhinal cortex neurons in male and female rodents (Arsenault et al., 2020)(see a review by (Ungar et al., 2014)). These morphological and physiological variations within different age and gender groups consequently yield divergent impacts on PI performance, a function involving the entorhinal cortex (Powell et al., 2021)(a Meta-analysis by (Farrer et al., 1997)). Beyond the entorhinal cortex, various other factors such as neurofibrillary tangle (NFT) formation (Ghebremedhin et al., 2001), cerebrospinal fluid (CSF) composition (Altmann et al., 2014), amyloid fibril levels (Cacciaglia et al., 2022), white matter microstructure (Srisaikaew et al., 2023), and functional connectivity within the default mode network (DMN) (Damoiseaux et al., 2012), are similarly influenced by age, gender, and APOE4. Consequently, they could also potentially influence the performance of path integration tasks. Behavioral studies in the past have yielded inconsistent findings regarding the roles of age, gender, and genotype in cognitive functions (Corder et al., 2004; Stevens et al., 2014; Wisdom et al., 2011; Zokaei et al., 2020). At the moment, the specific impacts of these factors on PI performance remain unresolved.

In the present study, we examined a group characterized by high-risk genes for AD development using a virtual reality-based path integration task (Fig. 1a,b). Within each trial of the experiment, participants experienced either instantaneous relocation (teleportation condition) or guided movement to a new location through visual cues (continuous navigation condition). Subsequently, they were tasked with navigating back to the initial starting point. Participants were encouraged to return to the starting location as accurate and quick as possible. Additionally, as part of a control task, participants were instructed to recollect the visual appearance of an image they had observed at the starting point (Fig. 1c). Our goal was to investigate how age, gender, the presence of high-risk AD genes, and continuous navigation influenced performance in the path integration task.

**Fig. 1.**
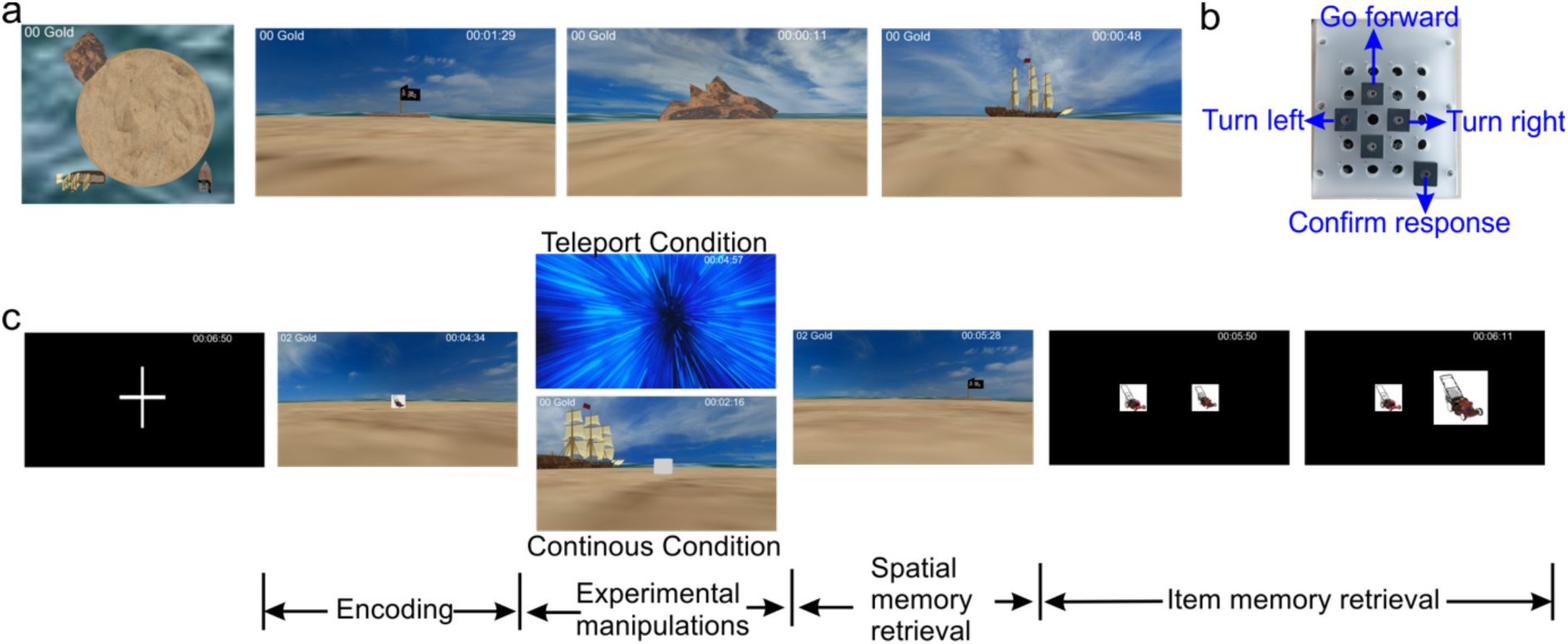
Experimental setup. **a**) Depiction of the island and its landmarks, presented from both an overhead perspective and a first-person viewpoint. **b**) Picture of the custom-made response box. It is made from a number pad by covering all buttons except arrow buttons and the enter button. **c**) The procedure of the experiment. Prior to each trial, a black screen with a white cross at the center was displayed for a jittered duration, ranging from one to two seconds. During the encoding phase, an image appeared on the island and subjects collected it by navigating to its location. Subjects were encouraged to remember details about the image and its location. Subsequently, their spatial memory was assessed. They were either teleported to a new location or directed to navigate continuously to a new location indicated by an empty cube. From the new location, they navigated to the target location. Following the spatial memory test, their item memory was evaluated. In this phase, both the target image and a highly similar lure image were presented on the monitor. Subjects chose the image that they had seen during the encoding phase. They were encouraged to perform the task as accurate and quick as possible.

## Results

### Demographic data

We assessed a group of 143 participants (99 females; mean age: 44.6±16.8 years), with 132 of them successfully genotyped. Among the genotyped participants, 37 carried at least one APOE e4 allele (Fig. 2a; Table 1). The proportion of APOE4 carriers observed in this study (25.87%) exceeded the previously reported ratio (∼15%; (Farrer et al., 1997); Binomial test; p = 0.00011). Subsequent data analyses focused on 31 individuals bearing APOE e3/e4 alleles (“carriers”) and 74 individuals with APOE e3/e3 alleles (“controls”). These two groups were matched for age (carriers: 43.1±17.1; controls: 45.3±15.9; t(103) = 0.62; p = 0.54; two-sample T test) and gender (carriers: 20/31 females; controls: 56/74 females; p = 0.15; Binomial analysis; Fig. 2b,c). Among female carriers and male carriers, the age match was evident (t(29)=0.14; p=0.44; two-sample T test; Fig. 2d), whereas male non-carriers were older than their female counterparts (t(72)=2.51; p=0.0060; two-sample T test; Fig. 2e). Additionally, target locations and the start searching locations were comparable between carriers and control groups (ps>0.05; Fig 1.f).

**Fig. 2.**
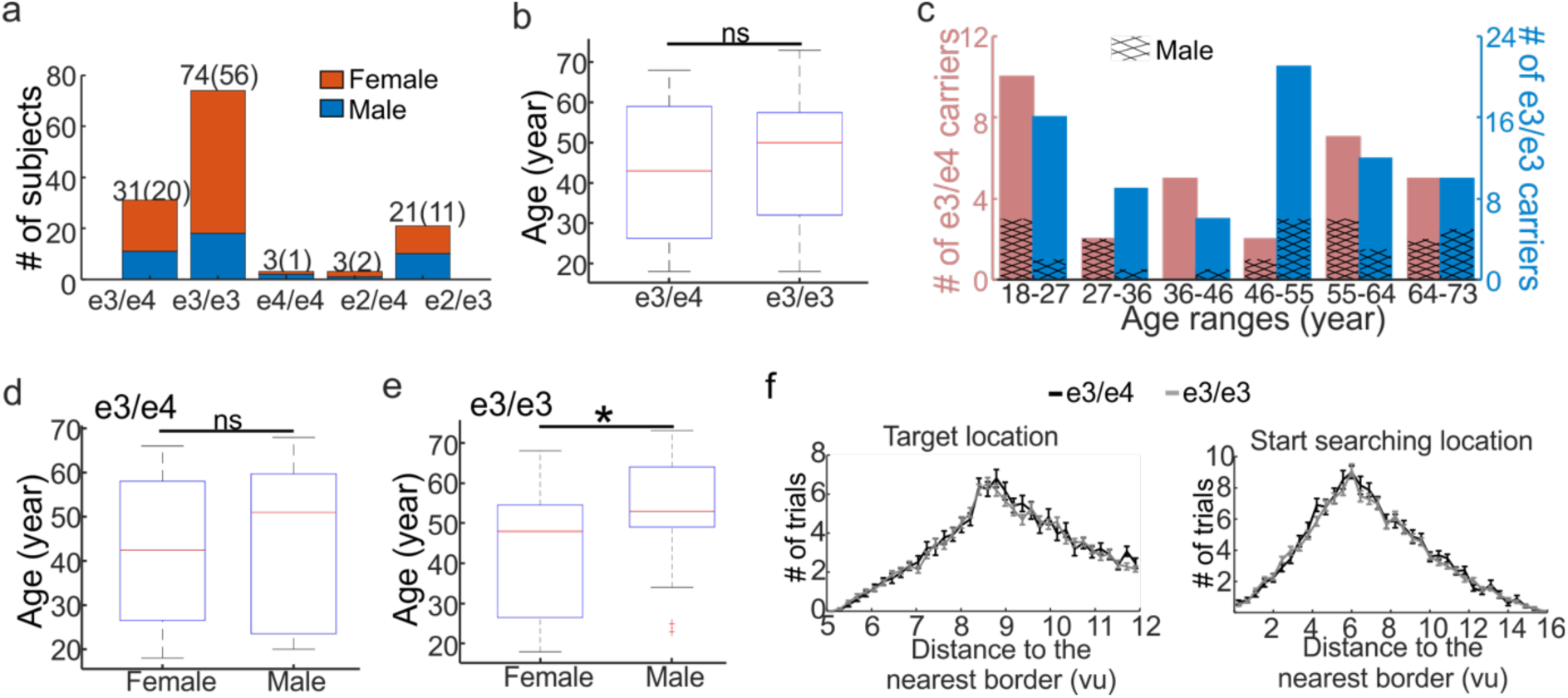
Demographic information of subjects. **a**) Distribution of subjects by gender and genotype carriers. The number displayed above each bar represents the total number of subjects, with the number in parentheses indicating the number of female subjects for each genotype. **b**) No noticeable difference was observed between carriers of high-risk genes (APOE e3/e4) and control subjects (APOE e3/e3). Ns: no significant difference. **c**) Representation of the age distribution, with high-risk gene carriers (e3/e4) depicted in red and control subjects (e3/e3) in blue. **d**) No age difference between female and male high-risk gene carriers. Ns: no significant difference. **e**) The age of male control subjects exceeded that of female control subjects. *: p<0.05. **f**) The allocation of the target and start searching locations was balanced between carriers and controls.

**Table 1.** Age information of subjects.

|  | APOE 3/4 | APOE 3/3 | APOE 4/4 | APOE 2/4 | APOE 2/3 |
| --- | --- | --- | --- | --- | --- |
| Age (mean±std) | 43.1±17.4 | 45.9±15.7 | 57.3±22.0 | 70.7±1.5 | 43.9±15.7 |

### Decomposing PI performance into translation and direction estimation

During navigation, subjects update their current location based on their knowledge of the translation and direction information (Benhamou & Séguinot, 1995; Fujita et al., 1990; Harootonian et al., 2020; Klatzky et al., 1990; Maurer & Séguinot, 1995). We thus examined how the accuracy of translation (i.e., distance error) and direction (i.e., angle error) estimation impacts the PI performance (i.e., the drop error; Fig. 2a). To this end, we first assessed the intraclass correlation coefficient (ICC = 0.25), showing that a substantial portion of the variation in the path integration performance was attributable to inter-subject differences (Fig. 2b). We then computed the Design EFFect (DEFF) to demonstrate the necessity of interpreting the dataset as a multilevel sample instead of a simple random sample. Our findings indicated that the sampling variance was approximately 27 times higher (DEFF=27.09>2) than if all subjects were part of a simple random sample. In light of this, we executed a multilevel linear regression analysis to model how translation and direction estimations performance contribute to path integration performance, accounting for variations at the subject level. The results of our analysis unveiled that both distance error and angle error exerted significant influence on drop error (distance error: B=0.68, SE=0.0054, 95%CI=0.67 to 0.69, p < 0.0001; angle error: B=0.070, SE=0.00057, 95%CI=0.069 to 0.071, p < 0.0001). Remarkably, distance error and angle error collectively accounted for 74% of the variance in drop error (R^2^ = 0.74). These findings corroborate the established findings (Benhamou & Séguinot, 1995; Fujita et al., 1990; Harootonian et al., 2020; Klatzky et al., 1990; Maurer & Séguinot, 1995) suggesting that path integration performance can be deconstructed into the accuracy of translation and direction estimation.

### Various factors impacting PI performance

Employing a multilevel linear mixed regression approach, we investigated how different factors—age, genotype, gender, and experimental conditions—along with their potential interactions, influence PI performance (i.e., the drop error). Within each subject, the experimental condition acted as a predictor, while age, gender, and genotype were considered predictors at the subject level. Our results revealed a significant interaction effect between gender and genotype in predicting the drop error (B=8.85, SE=2.78, 95%CI=3.35 to 14.34, p=0.0019; Fig. 2c). Further analyses unveiled that among APOE e3/e4 carriers, male subjects exhibited a larger drop error compared to female subjects (B=2.39, SE=0.79, p=0.0024), whereas in APOE e3/e3 carriers, the drop error was smaller in male subjects than in female subjects (B=1.40, SE=0.30, p=0.0052). In male subjects, APOE e3/e4 carriers showed a marginally larger drop error than APOE e3/e3 carriers (B=1.39, SE=0.72, p=0.054). In contrast, female subjects exhibited a reversal of this pattern, with APOE e3/e4 carriers showing a smaller drop error than APOE e3/e3 carriers (B=2.40, SE=0.59, p=0.0001). These findings suggest that the high-risk AD gene has a negative impact on PI performance in males but a positive influence in females. Furthermore, an interaction effect emerged between genotype and age (B=0.21, SE=0.098, 95%CI=0.013 to 0.40, p=0.038; Fig. 2d), indicating that the drop error increased more rapidly in APOE e3/e3 carriers compared to APOE e3/e4 carriers as age advanced. Subsequent analyses categorized subjects into young and old age groups (median age: 49 years) and found that the drop error was greater in the older age group compared to the younger group, regardless of genotype (APOE e3/e4: B=3.19, SE=0.77, p < 0.0001; APOE e3/e3: B=3.71, SE=0.60, p < 0.0001). No significant differences were observed between APOE e3/e4 carriers and APOE e3/e3 carriers within either age group (young group: B=0.56, SE=0.73, p=0.45; old group: B=0.37, SE=0.65, p=0.65). A trend toward a three-way interaction effect of age, gender, and genotype on drop error was also noted (B=-0.11, SE=0.056, 95%CI=-0.22 to 0.0022, p=0.055; Fig. 3e). Subsequent analyses indicated that over age, APOE e3/e3 carriers demonstrated a numerically faster increase in drop error compared to APOE e3/e4 carriers in female subjects (B = -0.0098, SE = 0.031, 95%CI=-0.070 to 0.051, p=0.75). Conversely, in male subjects, APOE e3/e4 carriers exhibited a faster increase in drop error than APOE e3/e3 carriers with age (B = 0.097, SE = 0.050, 95%CI=-0.00069 to 0.20, p=0.052). Although the experimental condition did not interact with other factors, it showed a significant main effect (B=7.98, SE=3.48, 95%CI=0.96 to 12.52, p=0.022; Fig. 3f), indicating that drop error was higher in the teleportation condition compared to the continuous navigation condition. This suggests that continuous navigation aids PI performance (Wiener et al., 2011). Given the decomposition of PI performance into translation and direction estimation, we further investigated how these different factors impact PI performance by affecting translation and direction estimation performance through mediation analysis (see Methods: Mediation analysis).

**Fig. 3.**
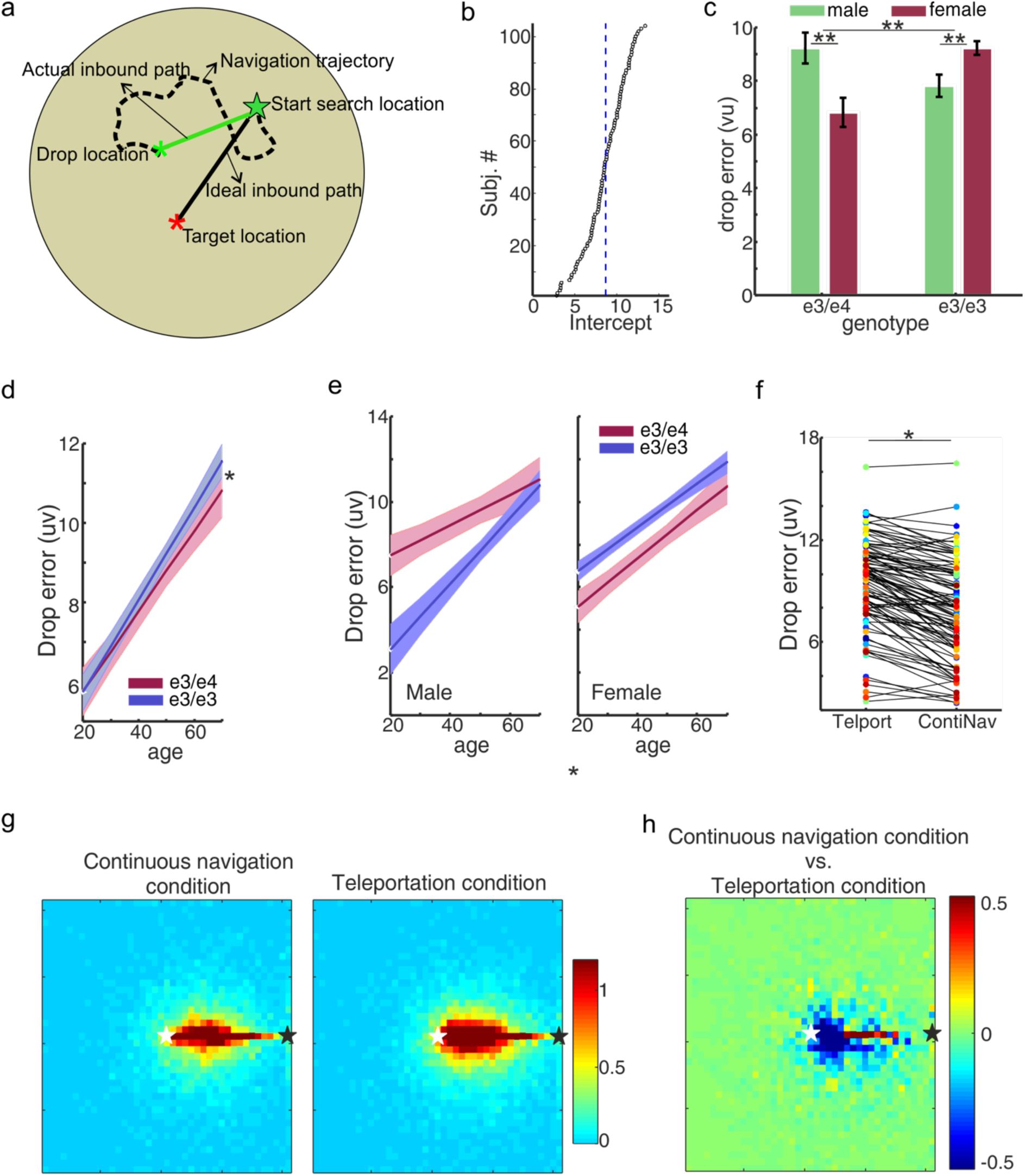
PI performance is influenced by multiple factors. **a**) Illustration of main terminologies. **b**) The PI performance varied over subjects. The dashed blue line indicates the ground mean. **c**) The interaction of genotype and gender influenced the PI performance. **d**) The interaction of genotype and age influenced the PI performance. **e**) The PI performance of the continuous navigation condition was better than that of the teleportation condition. **f**) A trend of the three-way interaction effect of genotype, gender, and age in influencing the PI performance. **g,h**) the difference in the navigation pattern between the continuous navigation condition and the teleportation condition. The hotter the color, the greater the number of visits to a specific location. White stars indicate the start searching location and black stars indicate the target location. *: p<0.05; **: p<0.01.

### The interaction of age and genotype influencing PI performance through the mediation of translation estimation performance

Our initial focus was on exploring how the interaction between age and genotype influences the drop error, with translation estimation serving as a mediator. To conduct this analysis, we employed a four-step mediation approach involving multilevel linear regression within each step (see Methods: Mediation analysis). The outcome of the first step, which highlighted the impact of the age-genotype interaction on the drop error, has been reported in the preceding section. Subsequently, we calculated that the age-genotype interaction also had a notable effect on the distance error (B=0.17, SE=0.077, 95%CI=0.019 to 0.33, p=0.028; step 2). Furthermore, the distance error served as a significant predictor of the drop error (B=0.74, SE=0.0078, 95%CI=0.72 to 0.75, p<0.0001; step 3). After accounting for the influence of the distance error, the interaction between genotype and age no longer predicted the drop error (B=0.082, SE=0.070, 95%CI=-0.056 to 0.22, p=0.24; step 4). These outcomes collectively indicate that the interaction of genotype and age exerts its effect on path integration performance primarily through the mediation of translation estimation performance. To quantify the mediation effect, we calculated the empirical mediator effect of the distance error and found it to be larger than all permuted values, resulting in a p-value < 0.0001 (see Methods).

We applied the same analysis to examine whether direction estimation also acted as a mediator in the impact of the age-genotype interaction on the drop error. However, the angle error failed to meet the criteria for mediation, as the interaction of genotype and age did not significantly affect angle error (in step 2: B=0.85, SE=0.65, 95%CI=-0.43 to 2.13, p=0.19). These findings underscore that the interplay between age and genotype, influencing path integration performance, is predominantly mediated by the accuracy of translation estimation. The summary of this section is illustrated in Fig. 5.

### The interaction of genotype and gender impacting PI performance, with mediation occurring through the performance of both translation and direction estimation

We further explored whether the interaction between genotype and gender, in predicting the drop error, was mediated by translation estimation. Our investigation revealed an interaction effect of genotype and gender on distance error (B=5.44, SE=2.21, 95%CI=1.07 to 9.80, p = 0.015; step 2). Subsequently, the distance error demonstrated a significant predictive role for the drop error (B=0.74, SE=0.0078, 95%CI=0.72 to 0.75, p<0.0001; step 3). Remarkably, even after accounting for the influence of the distance error, the interaction between genotype and gender continued to predict the drop error (B=4.94, SE=1.99, 95%CI=1.12 to 8.87, p=0.014; step 4). This suggests that the impact of the interaction between genotype and gender on path integration performance was partially mediated by translation estimation performance. Similarly, we applied a parallel analysis to angle error and found that the performance of direction estimation also acted as a partial mediator for the interaction effect of genotype and gender on path integration performance (step 2: B=45.77, SE=18.49, 95%CI=-9.23 to 82.31, p = 0.015; step 3: B=0.076, SE=0.00082, 95%CI=0.074 to 0.078, p<0.0001; step 4: B=5.42, SE=1.91, 95%CI=1.65 to 9.19, p=0.0053). Additionally, when simultaneously regressing out the effects of both distance and angle errors, the interaction between genotype and gender could no longer predict the drop error (B=2.01, SE=1.06, 95%CI=-0.088 to 4.11, p=0.061). Notably, all mediator effects proved statistically significant (ps<0.001), highlighting that the interaction effect of genotype and gender on path integration performance was jointly mediated by the performance of both translation and direction estimation. The summary of this section is illustrated in Fig. 5.

### The impact of elapsed searching time and searching path length on path integration performance depending on the experimental manipulation

As noted earlier, the continuous navigation condition exhibited superior path integration performance compared to the teleportation condition. We delved deeper into the contrast between these conditions and discovered that the elapsed searching time in the teleportation condition was longer than that in the continuous navigation condition (B = -1.95, SE=0.33, 95%CI=-2.61 to -1.29, p<0.0001). We found a similar result after excluding the non-moving periods (B = -1.32, SE=0.14, 95%CI=-1.60 to -1.04, p<0.0001). However, there was no significant difference in the elapsed searching path length between the two conditions (B = -1.00, SE=0.53, 95%CI=-2.05 to 0.48, p=0.062). Instead, the navigation trajectory was more deviated from the direct path for the teleportation condition than the continuous navigation condition (B=-5.27, SE=0.54, 95%CI=-6.33 to -4.21, p<0.0001; Fig. 3g, h). Further analysis within different experimental conditions unveiled that the drop error increased with the elapsed searching time in the teleportation condition (B = 0.029, SE=0.0060, 95%CI=0.017 to 0.041, p<0.0001), whereas no such correlation was evident in the continuous navigation condition (B = -0.0029, SE=0.0059, 95%CI=-0.014 to 0.0086, p=0.62). Conversely, the drop error decreased with the actual path length in the continuous navigation condition (B = -0.069, SE=0.0059, 95%CI=-0.080 to -0.058, p<0.0001), whereas this trend was not observed in the teleportation condition (B = -0.0076, SE=0.0061, 95%CI=-0.019 to 0.0043, p=0.21).

We proceeded to conduct a more detailed analysis within each experimental condition to explore whether the performance of translation and direction estimation mediated the influence of elapsed searching time and path length on the drop error. Our findings indicated that, in the continuous navigation condition, the searching path length had a partially mediated impact on the drop error through both the distance error (step 2: B = -0.011, SE=0.0046, 95%CI=-0.12 to -0.11, p<0.0001; step 3: B = 0.74, SE=0.010, 95%CI=0.72 to 0.76, p<0.0001; step 4: B = 0.015, SE=0.0043, 95%CI=0.0070 to 0.024, p=0.0070) and the angle error (step 2: B = -0.52, SE=0.044, 95%CI=-0.61 to -0.43, p<0.0001; step 3: B = 0.073, SE=0.0012, 95%CI=0.070 to 0.075, p<0.0001; step 4: B = -0.033, SE=0.0044, 95%CI=-0.041 to -0.024, p<0.0001). In contrast, in the teleportation condition, the elapsed searching time exhibited a partially mediated impact on the drop error through the angle error (step 2: B = 0.11, SE=0.048, 95%CI=0.013 to 0.20, p=0.025; step 3: B = 0.078, SE=0.0011, 95%CI=0.076 to 0.080, p<0.0001; step 4: B = 0.021, SE=0.0046, 95%CI=0.012 to 0.030, p<0.0001), while no such mediation was observed through the distance error (step 2: B = -0.0069, SE=0.0047, 95%CI=-0.016 to 0.0024, p=0.15). In summary, our findings indicate that elapsed searching time negatively impacts the performance in the teleportation condition partially mediated by direction estimation, while elapsed searching path length positively impacts the performance in the continuous navigation condition that is partially mediated by translation and direction estimation. The summary of this section is illustrated in Fig. 5.

### APOE4 gene impacting translation under-estimation during PI task

Our investigation into translation and direction estimation revealed that the actual inbound path length was shorter than the ideal inbound path length, indicating a signed distance error below zero (t(103) = 14.84, p<0.001). This underestimation phenomenon aligns with previous research (Fujita et al., 1990; Harootonian et al., 2020; Klatzky et al., 1999). Research has also established that subjects tend to exhibit regression responses, reflecting an inclination to underestimate longer distances while overestimating shorter distances based on their prior experiential estimation of the distance range (Petzschner & Glasauer, 2011; Teghtsoonian & Teghtsoonian, 1978). To further explore this, we conducted a median split of the trials. Our hypothesis was that subjects estimated the average distance during the first half and then applied regression responses in the second half. As a result, any underestimation effect observed in the first half trials would be invalid in the second half, as any underestimations would likely be counteracted by overestimations. However, our findings revealed that the signed distance error remained below zero for both halves (ts > 10.75, ps<0.001), and there was no significant difference in distance error between the two halves (t(103)=0.36, p=0.72). This indicates that our results do not support the repression response theory.

Interestingly, we observed that APOE4 carriers exhibited a larger signed distance error compared to controls (B=-1.34, SE=0.56, 95%CI=-2.45 to -0.24, p=0.018; Fig. 4a,b), suggesting that controls are worse in estimating distances. Conversely, no significant difference in angle error was found between carriers and controls (B = 2.09, SE=3.17, 95%CI=-4.19 to 8.37, p=0.51). These outcomes indicate that APOE4 carriers tend to underestimate distances more than their healthy counterparts. It’s worth noting that previous research has reported that APOE4 carriers tend to navigate along the boundary of the scenario (Coughlan et al., 2019; Kunz et al., 2015). We also calculated this aspect and did not find discernible difference between carriers and controls of their navigation pattern (ts < 0.58; ps > 0.55; Fig. 4 c,d).

**Fig. 4.**
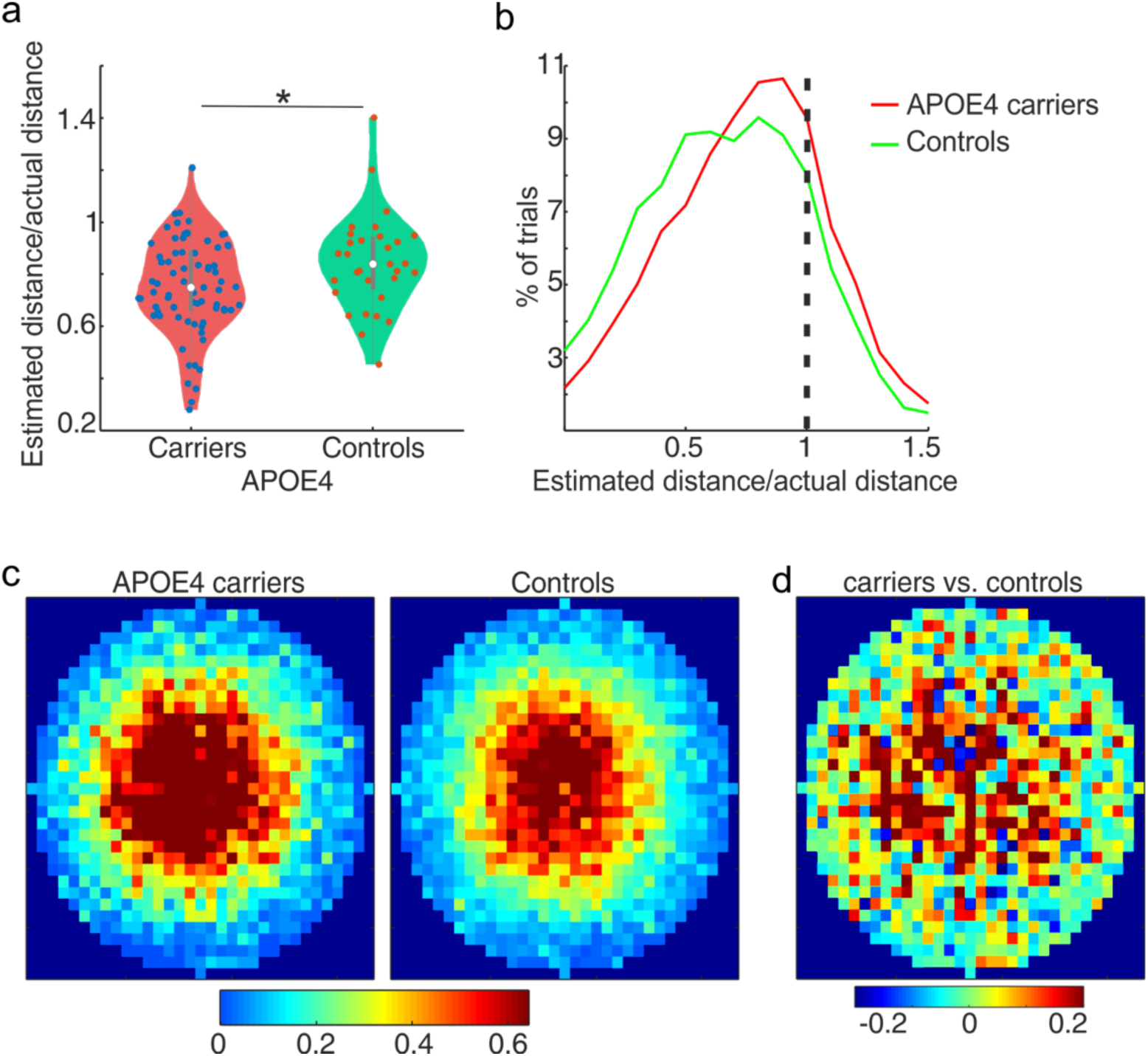
Difference between APOE4 carriers and control. **a,b**) APOE4 carriers underestimated less of the distance than controls (Bechtold et al., 2021). *: p<0.05. **c,d**) The navigation pattern of APOE4 carriers and controls did not differ.

**Fig. 5.**
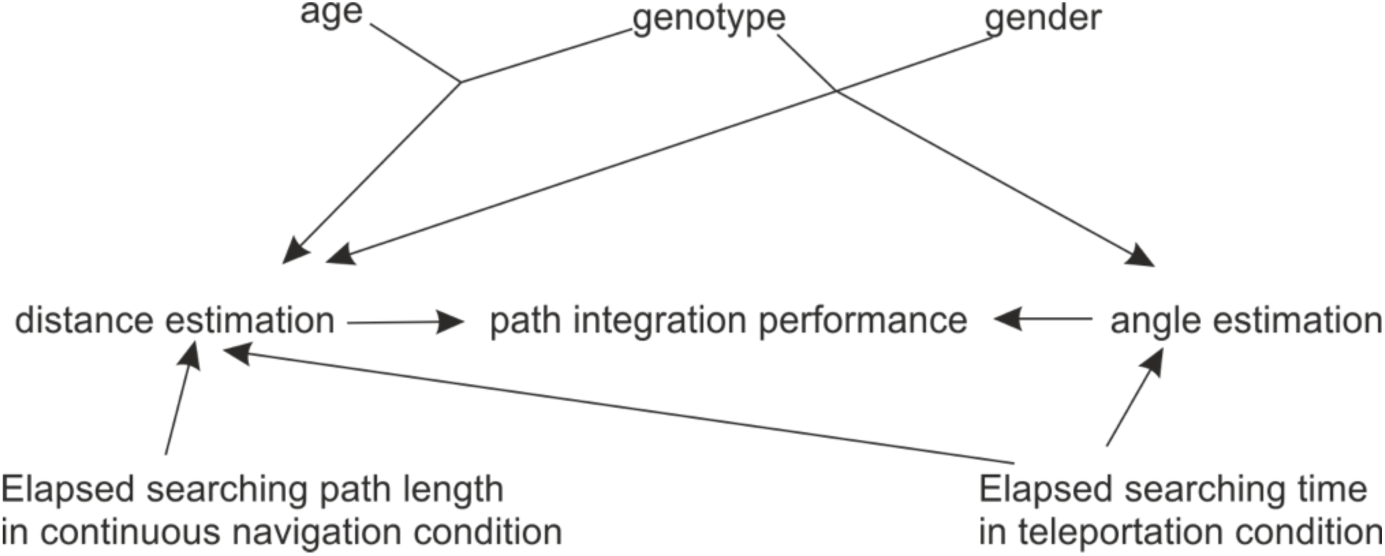
How various factors influence path integration (PI) performance through their effects on distance and angle estimation.

## Materials and methods

### Participants

One hundred and forty-three subjects (99 females; 44.6±16.8 years old) were recruited from the general population of the Ruhr area, either for monetary compensations or earning credit points for their participation. Subjects were free of neurological deficits and had no history of psychiatric disorders. All had a normal or corrected-to-normal vision and gave written informed consent before participating the study. Saliva samples were collected from subjects to analyze their APOE genotypes. DNA was extracted from saliva as described elsewhere (Miller et al., 1988). Using primers APOE_forward 5’-ACGGCTGTCCAAGGAG-3’ and APOE_reverse 5’-GGCCTGGTACACTGCC-3’, a 220 bp PCR product was generated containing both polymorphic regions, rs7412C/T and rs429358T/C. PCR was performed with 25 ng DNA, 300 nM primer each, and using GoTaq Green Master Mix (Promega, Germany) at an annealing temperature of 65.5°C in a final volume of 25 µl. Both polymorphisms were genotyped by RFLP-analysis-rs7412 using HaeII cutting the C-allele (220bp/202bp/18bp) and rs429358 using AflIII digest, cutting the T-allele (220bp/163bp/57bp). Out of the recruited individuals, 131 were successfully genotyped, and their demographic information were outlined in Table 1. Among the genotyped subjects, 31 were carriers of APOE e3/e4 alleles (“carriers”), while 73 carried APOE e3/e3 alleles (“controls”).

### Experimental stimuli

We built a virtual scenario (Panda3d; the Walt Disney Company) that was presented on a gaming laptop. The virtual scenario comprised a circular island with a radius of 16 virtual units, along with three distinctive landmarks surrounding the island: a small mountain, a small raft, and a pirate ship (Fig. 1a). A collection of 143 pairs of images was selected from the MST data pool (https://faculty.sites.uci.edu/starklab/mnemonic-similarity-task-mst/). These pairs were selected because they had been correctly labeled by a deep neural network (DNN) known as the Alexnet. Within each image pair, one image was randomly selected as the target stimulus, while the other one was designated as the lure stimulus. It’s important to note that the target and lure images in each pair were highly similar to each other.

We tested subjects’ spatial memory and item memory during the experiment (**Fig. 1c,d**). The spatial memory task encompassed two distinct conditions: the continuous navigation condition and the teleportation condition. In the continuous navigating condition, subjects navigated the virtual scenario to collect the target image (i.e., target location). Subsequently, they navigated to a new location indicated by a white cube (i.e., start searching location). From this point, they were instructed to navigate to the target location. Conversely, the teleportation condition involved subjects being instantly transported to a new location (i.e., start searching location) after collecting the target image. Following teleportation, they were instructed to navigate to the target location. Using a custom-designed number pad (**Fig 2b**), subjects utilized arrow buttons to navigate within the virtual scenario. The forward button enabled forward movement, the left and right buttons enabled turning, and the Enter button confirmed their responses upon reaching the target location (i.e., drop location). At the end of the spatial memory task, subjects received feedback indicated by the number of gold coins earned. Earnings amounted to three gold coins if the drop location lay within 4 virtual units of the target location, two coins within 8 virtual units, and one coin within 12 virtual units. The two spatial memory conditions were arranged alternately in a block design, with 10 trials in each block. Text was displayed on the monitor prompting subjects to switch experimental conditions at the beginning of each block. Notably, the target locations were more than 5 virtual units from the nearest border of the island and landmarks (Fig. 2e). It aimed to prevent subjects from relying on landmarks and borders to remember locations and instead to encourage them to perform metric computation (Philbeck et al., 2001).

In the item memory task, both target and lure images were displayed on the screen (**Fig. 2c**). Subjects utilized left and right arrow buttons to select one image, subsequently confirming their selections by pressing the Enter button. Accurate choices earned them one coin, while incorrect ones resulted in no coins being awarded.

### Experimental procedure

At the beginning of the experiment, subjects performed a practical session to familiarize themselves with the setup of the experiment. During the practical session, each block had one trial. Upon completing the practical session, subjects signaled the experimenter to initiate the actual experiment. The practical session followed the same procedure as the real experiment (Fig. 1c). At the beginning of each trial, a white cross appeared on the monitor for a jittered duration (1-2 seconds). Subsequently, subjects were initiated at a random location and directed towards the target image. Utilizing the number pad, they navigated towards the target image, which disappeared upon their arrival. Subjects were encouraged to memorize both the target image’s appearance and its location. Following this, they were either teleported to a new location (teleportation condition) or undertook continuous navigation to a new location marked by a white cube (continuous navigation condition). In the case of continuous navigation, the white cube disappeared once they reached it. Subsequently, subjects were instructed to navigate to the target location using the arrow buttons and confirm their response by pressing the enter button when reached the target location. Feedback was given based on the distance between the drop location and the target location, with greater accuracy earning more coins. Following the spatial memory’s feedback, two images were displayed on the monitor. Subjects employed the left or right arrow button to select the target image and pressed the Enter button to confirm their choice. Subsequently, they received feedback concerning their performance in the item memory task.

### Data analysis

We first clarified key terminologies employed in this paper.

*Target location:* The specific position of the target image.

*Drop location:* The location where subjects estimated the target image’s location.

*Start searching location:* The point to which subjects were teleported to in the teleportation condition or where subjects encountered the lure object in the continuous navigating condition.

*Elapsed searching path length:* The length of the path traveled from the start searching location to the drop location.

Elapsed searching time: The time taken to traverse from the start searching location to the drop location.

*Drop error:* The Euclidean distance between the target location and the drop location, Serving as an estimate of spatial memory performance.

*Ideal inbound path:* The direct line connecting the start searching location and the target location. *Actual inbound path:* The direct line between the start searching location and the drop location. *Angle error:* The angle formed between the ideal inbound path and the actual inbound path.

*Distance error:* The disparity between the length of the actual inbound path and the ideal inbound path.

The number of trials differed among subjects, with no distinction observed between APOE4 carriers and control subjects. Nevertheless, excluding subjects with a smaller number of trials did not alter the outcomes. As a result, we included all subjects in the data analysis.

#### Multilevel linear regression analysis

Previous studies have reported various factors that influence spatial memory, encompassing genotype, age, gender, and diverse cognitive strategies. To investigate the impacts of these factors, we constructed a comprehensive model by employing a linear mixed model to compute the effects of distinct factors and their potential interactions on the drop error (equation 1.1-1.4). Within individual subjects, the experimental manipulation was the predictor. At the subject level, the predictors included age, gender, and genotype.

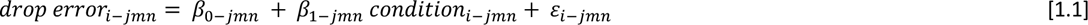

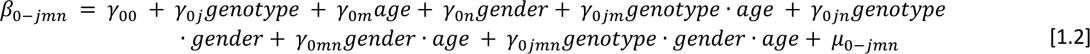

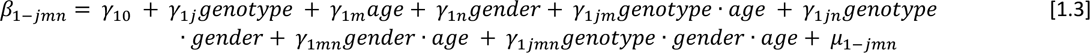

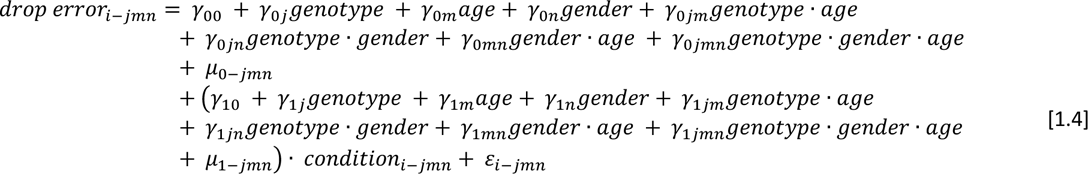

#### Mediation analysis

The mediation analysis followed a structured procedure consisting of four steps (Frazier et al., 2004; Judd & Kenny, 1981), with each step employing a linear mixed model. Step 1 showed the predictor variable (*var_predictor_*) predicted the response variable (*var_response_*; equation 2.1). Step 2 showed that the predictor variable predicted the mediator variable (*var_mediator_*; equation 2.2). Step 3 showed that the mediator variable predicted the response variable (equation 2.3); Step 4 established that the mediator variable mediated the relation between the predictor and response variables (equation 2.4). Meeting the criteria of the first three steps indicated that the association between the predictor and response variables was mediated by the mediator variable. Complete mediation was evident if the predictor variable had no direct effect on the response variable when including the mediator variable in Step 4. If a direct effect remained, it indicated partial mediation. The mediator effects were quantified (*effect_mediator_*; equation 2.5). The statistical significance of the *effect_mediator_* was determined through a permutation analysis with 10,000 iterations. In each iteration, the predictor variable was randomly shuffled, generating 10,000 permuted *effect_mediator_*s. The significance level of the empirical *effect_mediator_* was defined as its relative rank within these 10,000 permuted values. If the empirical *effect_mediator_* was larger than 95% of the permuted values, it was deemed statistically significant and indicated that the predictor variable influenced the response variable mediated by the mediator variable.

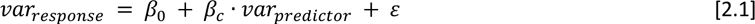

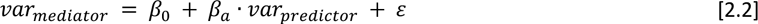

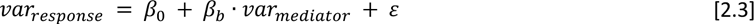

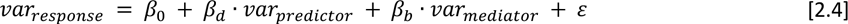

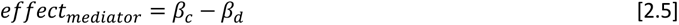

## Discussion

### The contribution of the entorhinal cortex to translation and direction estimation

During the execution of the path integration task, individuals monitor their translational and directional information to continuously update their spatial locations relative to the starting point. The spatial pattern exhibited by the firing field of grid cells makes them well-suited for facilitating distance estimation (Hafting et al., 2005). Research involving lesion studies in rodents has underscored the crucial function of the medial entorhinal cortex (MEC) in linear integration processes (Jacob et al., 2017) . Furthermore, empirical evidence has demonstrated that grid cells encode distance information as rodents navigate through a T-shaped maze (Kraus et al., 2015). In the context of human subjects, neuroimaging investigations have indicated that the activation of the entorhinal cortex (EC) is inversely correlated with the spatial proximity to the destination point during navigation (Spiers & Maguire, 2007). This discovery was corroborated by another study that established the EC’s involvement in encoding the Euclidean distance to the goal (Howard et al., 2014). Additionally, distortion of grid cell activity near boundaries also leads to diminished path integration performance (X. Chen et al., 2015). In recent times, the concept of grid-cell-like representations (GCLRs) has emerged from the analysis of BOLD signals or local field potentials (D. Chen et al., 2018; Doeller et al., 2010; Maidenbaum et al., 2018). It’s worth noting that GCLRs capture not just grid cell activity but also the interplay between grid cells and head-direction cells. This suggests their potential to provide both translation and direction information concurrently. Our data support the plausible role of EC in estimating distance and angles, as evidenced by the full mediation of the interaction effect between genotype and age on path integration performance through distance estimation. Likewise, distance and angle estimation jointly mediate the interaction effect of gender and genotype. Further studies are needed to tease apart the role of EC in distance and direction estimation during PI tasks.

### Elapsed path length is not necessary for inducing accumulated error in PI tasks

In our current study, we made an intriguing observation that the elapsed searching path length did not exert an influence on the drop errors in the teleportation condition. Furthermore, we noticed a reduction in drop errors as the elapsed searching path length increased under continuous navigation conditions. These findings diverge from prior research, which indicated that the PI performance decreased with accumulated navigation path length because the estimating error increased with the increased navigating path length (Lakshminarasimhan et al., 2018; Lappe et al., 2007, 2011). One plausible explanation for this inconsistency could be the presence of the landmarks. In our study, we deliberately designed the task to encourage subjects to engage in metric computation while searching the target location instead of relying on landmarks (Philbeck et al., 2001). However, we cannot entirely dismiss the possibility that subjects might have relied on landmarks to orient themselves within the environment. Landmarks, during navigation, serve to recalibrate the ongoing location, aligning with the theory proposing a non-uniform, wrapped cognitive map (Bruns & Chamberlain, 2019; Carbon & Hesslinger, 2013; Cartwright & Collett, 1987; Cheung et al., 2012; Collett & Graham, 2004; Etienne et al., 1996, 1998; Hirtle & Jonides, 1985; Moar & Bower, 1983; Omer, 2013; Ra et al., 2017). According to the theory, a cognitive map is not a uniform two-dimensional representation but rather a wrapped multi-dimensional map. Areas devoid of landmarks are encoded with compressed precision, while regions proximity to landmarks are represented in an expanded manner. This could potentially explain the phenomenon of distance underestimation in PI performance. It is important to note that we carefully controlled for running speed, a factor previously identified as contributing to increased errors in PI performance (Stangl et al., 2020) .

### Reduced underestimation of inbound distances in APOE4 carriers

Our findings revealed a prevalent tendency among subjects to underestimate inbound distances. This outcome aligns with prior research indicating that individuals often display a tendency to underestimate distances traveled over inbound paths during path integration tasks (Harootonian et al., 2020; Philbeck et al., 2001). To delve deeper, our study showed that individuals carrying the APOE4 gene variant demonstrated a higher accuracy in estimating distances. In contrast, those without the APOE4 gene variant exhibited a more pronounced tendency to underestimate distances. The seemingly counterintuitive nature of this observation could potentially be attributed to the notion that a reduced tendency to underestimate distances might serve as a distinct behavioral marker for cohorts carrying genes associated with a higher risk of developing Alzheimer’s disease (AD). The hippocampus and EC are among the first target regions during the pathological progression of AD. Previous investigations have illuminated that these regions exhibited heightened activity during cognitive tasks (Nuriel et al., 2017; Tran et al., 2016). This elevated neural activity has been posited as a compensatory mechanism to counteract the damage incurred during disease development. Consequently, cognitive functions reliant on these hyperactive regions are also bolstered. Empirical support for this hypothesis has been acquired from studies indicating that individuals carrying the APOE4 gene variant outperform their non-carrier counterparts in tasks involving short-term memory, though not in tasks involving long-term memory (Stevens et al., 2014; Wisdom et al., 2011; Zokaei et al., 2020) . From this standpoint, we postulate that in the early stages of pathological development, the hippocampus and the entorhinal cortex are targeted, prompting their hyperactivation. This enhanced neural response subsequently augments tasks like distance estimation that call upon these specific brain regions for processing.

### Differential effects of the APOE4 genes over gender groups

The expression of the APOE4 gene exhibits distinct patterns within various gender cohorts. In male subjects, the PI performance of individuals carrying the APOE4 gene was notably poorer compared to APOE4 non-carriers. This finding concurs with the proposition that the pathological trajectory of Alzheimer’s disease (AD) initiates early in life (Bateman et al., 2012; Perrin et al., 2009). Moreover, this outcome aligns with previous research demonstrating impaired working memory performance among male APOE4 carriers (Reinvang et al., 2010; Zokaei et al., 2017) . However, an intriguing reversal of effects emerged in female subjects, where the PI performance of APOE4 carriers surpassed that of APOE4 non-carriers. This result contrasts with the general trend observed in prior studies, which indicated a more pronounced negative impact of APOE4 on females compared to males (as reviewed in (Ungar et al., 2014)). Upon reviewing existing literature that outlines gender differences in APOE4 carriers (Corder et al., 2004; Damoiseaux et al., 2012; Fleisher et al., 2005; Ghebremedhin et al., 2001; Lehmann et al., 2006; Mortensen & Høgh, 2001; Reinvang et al., 2010; Swan et al., 2005), we observed that the heightened negative influence of APOE4 in females predominantly manifests in older groups, typically ranging between 60 to 75 years of age (comprising approximately 9,000 subjects in total). As a result, we postulate that the divergent effect observed in our current study could be attributed to the relatively youthful age distribution of our cohort. Indeed, we identified a trend suggesting a three-way interaction effect between age, gender, and genotype. To comprehensively elucidate the impact of APOE4 on different age groups within both male and female populations, additional investigations are imperative.

## Conclusions

Employing a virtual navigation paradigm, we investigated various factors influencing path integration performance. Our observations revealed that the interaction of genotype and age impacted the path integration performance mediated through distance estimation. Moreover, the interaction of genotype and gender influenced path integration capabilities mediated by the distance and angle estimation. Additionally, under conditions of continuous navigation, the elapsed searching path length predicted the path integration performance partially mediated by distance and angle estimation. In contrast, in the teleportation condition, the elapsed searching time predicted the path integration performance partially mediated by the angle estimation.

## Acknowledgement

We would like to thank all students who were involved in data collection. They are Deirdre Fiona Fels, Jan Alexander Heistermann, Alica Jungnickel, Sandra Ostrowski, and Hanna Sophie Schmidt.

## Notes

### Competing Interest Statement

The authors have declared no competing interest.

